# Phylogenetic trees of closely related bacterial species and subspecies based on frequencies of short nucleotide sequences

**DOI:** 10.1101/2022.05.10.491390

**Authors:** Yoshio Nakano, Yusaku Domon, Kenji Yamagishi

**Affiliations:** Nihon University School of Dentistry, Department of Chemistry, Tokyo, 101-0061, Japan; Nihon University, College of Engineering, Department of Chemical Biology and Applied Chemistry, Fukushima, 963-8642, Japan

## Abstract

Bacterial phylogenetic analyses are commonly performed to explore relationships among various bacterial species and genera based on their 16S rRNA gene sequences; however, the results are limited by mosaicism, intragenomic heterogeneity, and difficulty in distinguishing between related species. In this study, we aimed to perform genome-wide comparisons of different bacterial species, namely *Escherichia coli* and *Shigella* and *Yersinia, Klebsiella*, and *Neisseria* spp. or serotypes of *Listeria monocytogenes*, based on their K-mer profiles to construct phylogenetic trees. Pentanucleotide frequency analysis (512 patterns of 5 nucleotides each) was performed to distinguish highly similar species from each other, such as *Yersinia* species, along with *Shigella* spp. and *E. coli*. Moreover, *Escherichia albertii* strains were clearly distinguished from *E. coli* and *Shigella*, despite being closely related to enterohemorrhagic *E. coli* in the phylogenetic tree. In addition, the phylogenetic tree of *Ipomoea* species based on pentamer frequency in chloroplast genomes correlated with previously reported morphological similarities. Furthermore, a support vector machine clearly classified *E. coli* and *Shigella* genomes based on pentanucleotide profiles. These results suggest that phylogenetic analysis based on penta- or hexamer profiles is a useful alternative for microbial phylogenetic studies. In addition, we introduced an R application, Phy5, which generates a phylogenetic tree based on genome-wide comparisons of pentamer profiles. The online version of Phy5 can be accessed at https://phy5.shinyapps.io/Phy5R/.

## Introduction

Currently, prokaryotic phylogenetic classification depends on 16S rRNA gene sequences, which are ubiquitously present and highly conserved in bacteria, but species with more than ≥ 99% identity based on 16S rDNA sequencing are rarely classified. In such cases, one or more additional conserved genes are often used as a secondary candidate for indexing during phylogenetic analysis, such as *gyrB* for *Shigella, Salmonella*, and *E. coli* [1]; *trpA, trpB, pabB*, and *putP* for *Shigella, Salmonella*, and *E. coli* [2]; and *thrA, trpE, glnA, tmk*, and *dmsA* for *Yersinia* spp. [3]. Since these candidate genes are not ubiquitously present among different bacterial species, the selection of an appropriate candidate gene is challenging. Furthermore, phylogenetic classification based on a single gene sequence is not reliable because the selected gene may have undergone horizontal transfer among bacterial species and thus does not reflect the background or history of the species.

To avoid these defects in phylogenetic analysis depending on a single gene, multi-locus sequence analysis/typing (MLSA/MLST) or core genome multi-locus sequence typing (cgMLST) is applied. For *Yersinia*, a genotyping strategy based on five genes was first developed in 2005 [4], and a seven-gene MLST scheme was established to differentiate among the three human pathogenic species: *Y. pestis, Y. pseudotuberculosis*, and *Y. enterocolitica* [5]. However, these MLST systems were not applicable at the species level. A genus-wide seven-gene MLST [6] and core-genome MLST [7] was developed for phylogenetic analysis of *Yersinia* species, which had notably better resolution and phylogenetic precision than the previous MLST methods. The core-genome MLST scheme facilitates high-resolution and efficient identification of various pathogens [8–12]. MLST with only a few genes does not reflect the evolutionary history of whole genomes. Species-specific, fixed sets of conserved genes throughout the genome are potentially used by cgMLST.

Among the abovementioned examples, the genetic evolution of *Yersinia* spp. is poorly understood. *Yersinia* is a gram-negative bacterium consisting of 19 species and includes 3 prominent human pathogens: *Y. pestis, Y. pseudotuberculosis*, and *Y. enterocolitica* [13]. Various virulence genes from Yersinia species have been reported, and phylogenetic analysis has been performed based on the sequence homology of specific virulence genes selected from those genes. In this study, we aimed to develop a method to phylogenetically any species or subspecies, including non-virulent strains, independent of specific gene sequence homology to characterized species.

The K-mer frequency in DNA fragments represents a genome-specific parameter to analyze genome diversity [14, 15]. The self-organizing map (SOM) based on K-mer nucleotide frequency analysis is used to cluster and visualize DNA fragments derived from closely related eukaryotes [16] or prokaryotes [17] in environmental samples. Sims et al. (2009) reported an alignment-free method, including the feature (or K-mer) frequency profiles (FFPs) of whole genomes [18], and the phylogenetic tree of 10 mammalian species was constructed from intron genomes based on 18-mer frequency profiles, and the resultant tree closely reflected the accepted evolutionary history. Furthermore, they reported a whole-genome-based phylogenetic tree of *E. coli* and *Shigella*, which was constructed using 24-mer frequency profiles [19]. All possible combinations of 24-mer fragments produced ca 8.4 million features. In this study, based on 512 pentamer combinations, we aimed to perform phylogenetic separation of *E. coli* and *Shigella* and *Yersinia, Klebsiella*, and *Neisseria* spp. or serotypes of *Listeria monocytogenes*.

A support vector machine (SVM) is a supervised machine-learning model and is one of the most recently developed classifiers. It segregates classes using hyperplanes generated by mapping the predictors into a new, higher-dimensional space (the feature space), wherein they can be linearly segregated. We expected that SVM would help classify numerous samples of species from genetic sequences in a short period based on K-mer frequencies and such an application is verified here.

## Materials and methods

### Genome Sequences

In total, 110 *Yersinia*, 888 *E. coli*, 92 *Shigella*, 280 *Campylobacter*, 561 *Klebsiella*, 67 *Listeria*, 188 *Neiserria*, and 18 *E. albertii* genome sequences were obtained from the ftp site at NCBI Microbial Genomes (ftp://ftp.ncbi.nlm.nih.gov/genomes/Bacteria/). Some strains contained ≥ two genomes or megaplasmids. Genomes or plasmids with molecular size more than one-tenth the size of the largest genome or plasmid were combined into one nucleotide sequence, and those with size less than one-tenth the size of the largest genome or plasmid were excluded.

The chloroplast genomes of *Ipomoea* species were obtained from the ftp site at NCBI plastid sequences (https://ftp.ncbi.nlm.nih.gov/refseq/release/plastid/).

### K-mer frequencies and phylogenetic trees

The tri-, tetra-, penta-, and hexanucleotide frequencies in bacterial genomes of each sample were determined using R 4.12 (http://www.r-project.org) with the Biostrings package. Degenerated frequencies of K-mer nucleotides, in which complementary K-mer pairs (e.g., AAA vs. TTT) were considered the same nucleotide string, were used to construct phylogenetic trees via hierarchal cluster analysis with Manhattan distance and Ward’s algorithm.

### Phylogenetic trees based on 16S rRNA gene sequences

We extracted 16S rDNA sequences from the aforementioned genome sequences and aligned them using MAFFT [20]. These sequences were used to construct neighbor-joining trees, using MUSCLE [21].

### Machine learning

Analysis and classification of the bacterial genome of each strain were accomplished using R 4.12 and the e1071 packages for the SVM with the radial basis function (RBF).The radial kernel function transformed the data using the non-linear function *k*(*x*1, *x*2) = *exp*(*γ| x*1 − *x*2|2), where *γ* determines the RBF width, unless otherwise specified. Classification through machine learning was performed using the leave-one-out cross-validation method, i.e., one sample was classified through supervised machine learning using the other 947 samples

### R application

Phy5 was coded in R and Bioconductor using the shiny, biostrings, ape, and pvclust packages. These can be further customized or extended using HTML and CSS as Shiny applications.

## Results and Discussion

Phylogenetic analysis based on the 16S rRNA gene sequences is not suitable for closely related species. A bacterial 16S rRNA gene sequence contains 9 variable regions, and the 500-nucleotide long hypervariable V1–V3 region or the 800-nucleotide long V1–V4 region is used for phylogenetic analysis. In closely related species, these variable regions do not vary sufficiently to perform phylogenetic analysis. *E. coli, Salmonella, Shigella, Yersinia*, and *Klebsiella* from the Enterobacteriaceae family share up to 99% sequence identity based on their full 16S rRNA gene sequences [22]. In many cases, bacterial operational taxonomic units (OTUs) are defined by 97% identity between each other, thus a set of species sharing 99% rRNA gene sequence identity cannot be differentiated using 16S rRNA gene sequencing. In addition, prokaryotic species have two types of rRNA operons, and horizontal gene transfer is observed among many bacterial species [23]. The genetic interoperability and promiscuity of 16S rRNA in the ribosomes of an extremely thermophilic bacterium, *Thermus thermophilus*, has been reported by Miyazaki et. al. (2019) and they suggested that horizontal gene transfer promoted adaptive evolution, that is the “random patch model” for ribosomal evolution [24].

Figure 1 shows the results of our proposed method for phylogenetic analysis of closely related species that are resistant to multicopy, horizontal transfer, or chimera-formation of the 16S rRNA genes and do not require species-specific gene sets, such as MLSA/MLST. Such a method will help perform rapid and easy phylogenetic analysis of any species. For example, when a microbiologist attempts interspecies crossing of morning glory, without being familiar with the genus, *Ipomoea*, the proposed method could provide rapid and useful information about the phylogenetic relations between a collection of species of the genus to study the genetic characteristics of MLSA/MLST. Whole-genome comparison is required if specific genes are not selected for phylogenetic analysis, and such a comparison would limit inconsistencies due to gene transfer, multicopy, or chimera-formation of the 16S rRNA genes. As mentioned above, SOM based on K-mer nucleotide frequency analysis has been reported for environmental samples [16, 17]. Frequency analysis of short nucleotide sequences is applicable for any genus or species, as it does not require the selection of specific genes for phylogenetic analysis, as shown in Fig. 1. In addition, complete genome sequences are not required for this method, and accumulations of short nucleotide sequences, such as the results of NGS, are sufficient for the analysis. Based on this idea, we demonstrate the efficiency and practicality of phylogenetic analysis by pentanucleotide frequencies.

**Fig 1.**
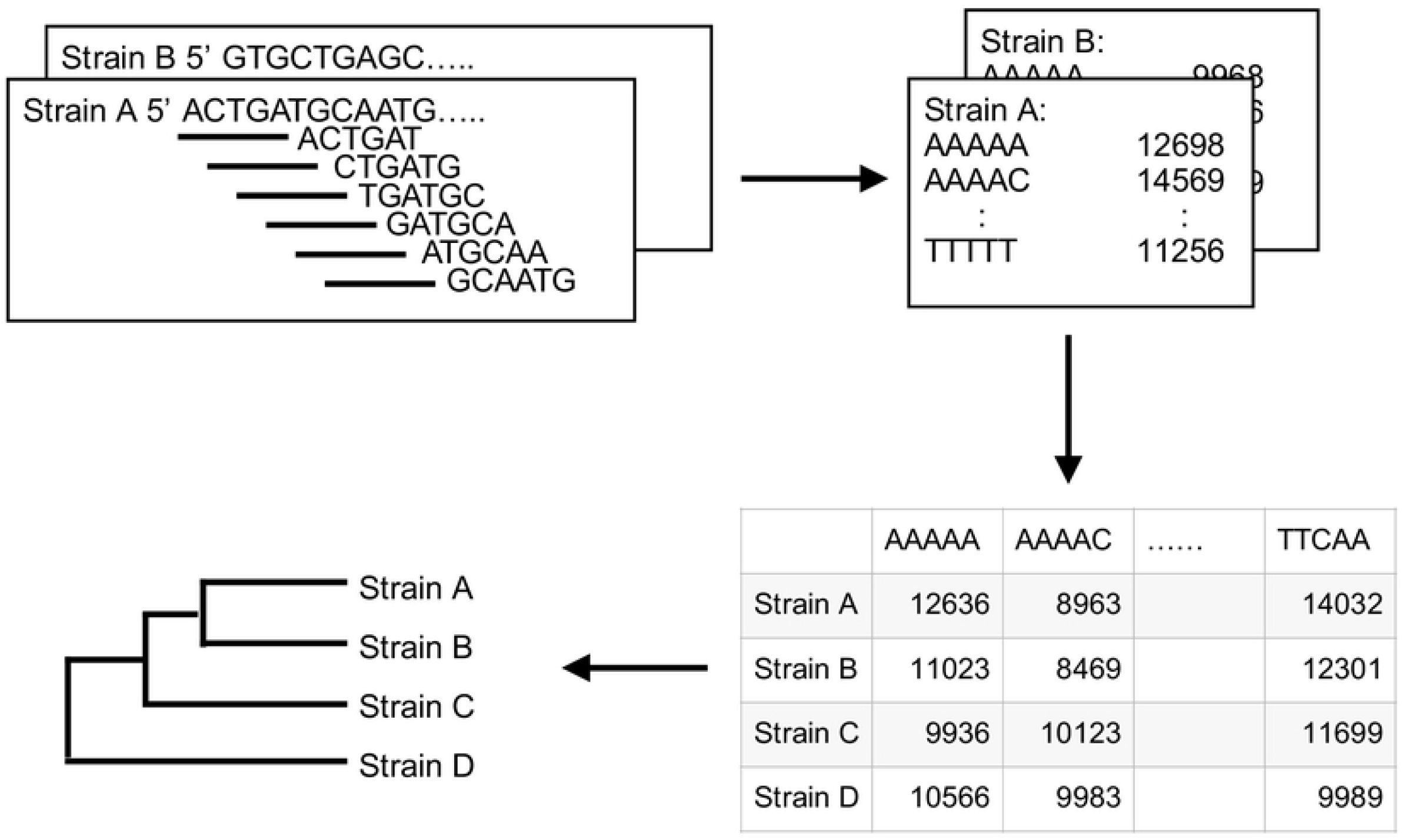
Scheme for constructing a phylogenetic tree based on pentanucleotide frequencies using Phy5.

Degenerated pentanucleotide frequencies were calculated from the DNA sequences of 110 *Yersinia* genomes (Supplementary Data 1), and in accordance with the results, phylogenetic analyses were performed (Fig. 2A). The phylogenetic tree, based on pentamer frequencies with 100% bootstrap values, demonstrated that *Y. pestis, Y. pseudotuberculosis, Y. enterocolitica*, and *Y. ruckeri* clearly separated from one another, as shown in Fig. 2A, whereas the analyses based on 16S rRNA gene sequences (Fig. 2B) could not distinguish among these species. Furthermore, phylogenetic trees based on tri-, tetra-, penta-, and hexa-nucleotide frequencies (Supplementary Data 1-4) were constructed (Supplementary Fig 2). The last three trees distinguished the three species of *Yersinia* with 100% bootstrap values. Hexamer frequencies distinguished all strains, with ≥ 99% bootstrap values. However, this result does not necessarily determine the minimum number of nucleotide frequencies required for phylogenetic tree analysis. Furthermore, this number cannot be calculated in advance because the diversity in DNA sequences is not constant across various species and genera. However, pentanucleotide frequencies are sufficient for phylogenetic analysis of *Yersinia* and other species described in this study.

**Fig 2.**
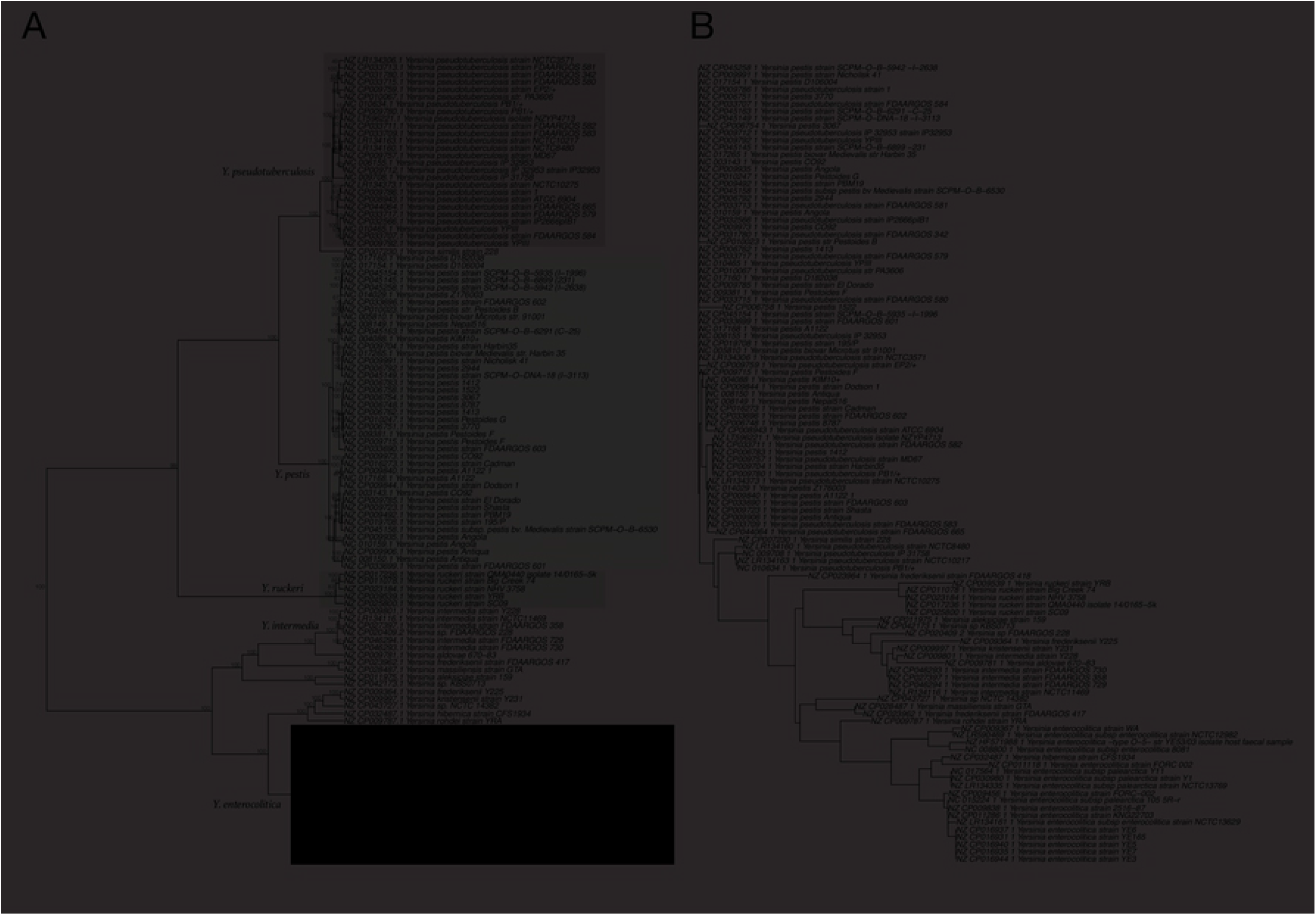
Phylogenetic trees of 110 Yersinia strains based on pentanucleotide frequencies (A) and 16S rRNA gene sequences (B). The trees were constructed using the Manhattan distance and Ward’s algorithm (A) or the neighbor-joining method (B). The numbers at the nodes indicate the percentage occurrences among 1,000 bootstrap values. Separated groups of species are highlighted.

Furthermore, the phylogenetic trees for *E. coli/Shigella* (Fig. 3, Supplementary Fig 2), *Campylobacter* (Supplementary Fig 3), *Klebsiella* (Supplementary Fig 4), *Listeria* (Supplementary Fig 5), *Neisseria* (Supplementary Fig 6), and *Escherichia albertii*, including *E. coli* and *Shigella* (Supplementary Fig 7) species/strains were generated using pentanucleotide frequencies (Supplementary Data 1).

**Fig 3.**
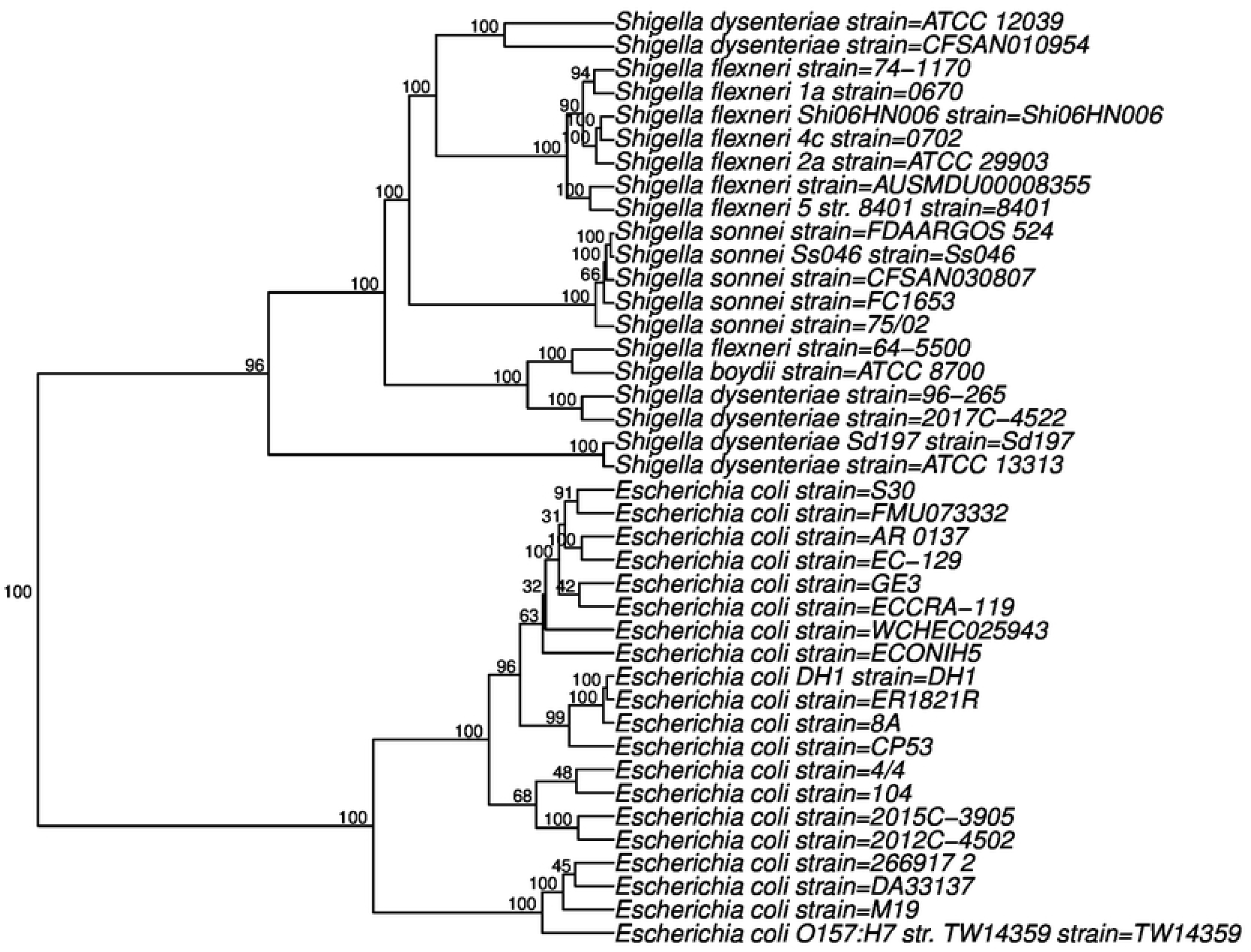
Phylogenetic trees of 20 random sequences from 888 sequences of *E. coli* and 20 random sequences from 92 sequences of *Shigella* based on the pentanucleotide frequencies.

*E. coli* and *Shigella* are difficult to distinguish on the basis of their 16S rDNA sequences alone, and numerous alternatives have been proposed for their phylogenetic analysis [1, 2, 25]. For example, Fukushima et al. reported phylogenetic analyses of

*Salmonella, Shigella*, and *E. coli* strains in accordance with the sequences of the *gyrB* gene [1], whereas Escobar-Páramo et al. used four chromosomal genes, (*trpA, trpB, pabB*, and *putP*), or three plasmid genes, (*ipaB, ipaD*, and *icsA*) [2]. Phylogenetic trees based on pentanucleotide frequencies for the four species of *Shigella* and *E. coli* strains are distinct (Supplementary Fig. 2), and bootstrap values indicate the significance of the separation.

*E. coli* and *Shigella* are difficult to distinguish on the basis of their 16S rDNA sequences alone, and numerous alternatives have been proposed for their phylogenetic analysis [1, 2, 25]. For example, Fukushima et al. reported phylogenetic analyses of *Salmonella, Shigella*, and *E. coli* strains in accordance with the sequences of the *gyrB* gene [1], whereas Escobar-Páramo et al. used four chromosomal genes, (*trpA, trpB, pabB*, and *putP*), or three plasmid genes, (*ipaB, ipaD*, and *icsA*) [2]. Phylogenetic trees based on pentanucleotide frequencies for the four species of *Shigella* and *E. coli* strains are distinct (Supplementary Fig. 2), and bootstrap values indicate the significance of the separation.

*E. albertii* was originally classified as a *Hafnia alvei*-like stain isolated from human stool samples in the early 1990s and was suspected of causing diarrhea [26]. *I* t has recently been recognized as a close relative of *E. coli*. Further, *E. albertii* strains were earlier found to be closely related to strains of *Shigella boydii* serotype 13, a distant relative of *E. coli*, representing a divergent lineage in the genus *Escherichia* [27]. Furthermore, Hyma et al. reported that the *E. albertii-Shigella* B13 lineage is estimated to have diverged from an *E. coli*-like ancestor 28 million years ago. Herein, we constructed the phylogenetic tree of *E. albertii, E. coli*, and *Shigella* from pentanucleotide frequencies (Fig. 4). Figure 4 shows the phylogenetic relationships of lineages in the three species. *E. albertii* strains and enterohemorrhagic *E. coli* strains, including O157, O121, and O111, are closely related to each other and are on separate branches distinct from *Shigella* and nonpathogenic *E. coli*, such as strain K-12. Figure 4 shows that *Shigella* strains are distinguished from enteroinvasive E. coli (EIEC), and these groups form a large branch separated from the other *E. coli* strains, including K12. Three clusters within *Shigella* appear to have diversified in a period of 35,000–270,000 years [25]. One *E. coli* and two *Shigella* species were identified in opposite genus (Supplementary Figure 3). *E. coli* NCTC11104 was classified as *Shigella*, whereas it is now identified as Citrobacter Sc16. *S. flexneri* C32 and *Shigella* sp. PAMC 28760 were classified as *E. coli*. The former bacterium was isolated from a Himantormia sp. lichen in Antarctica; thus, its genetic history could be different from that of other *Shigella* spp.

**Fig 4.**
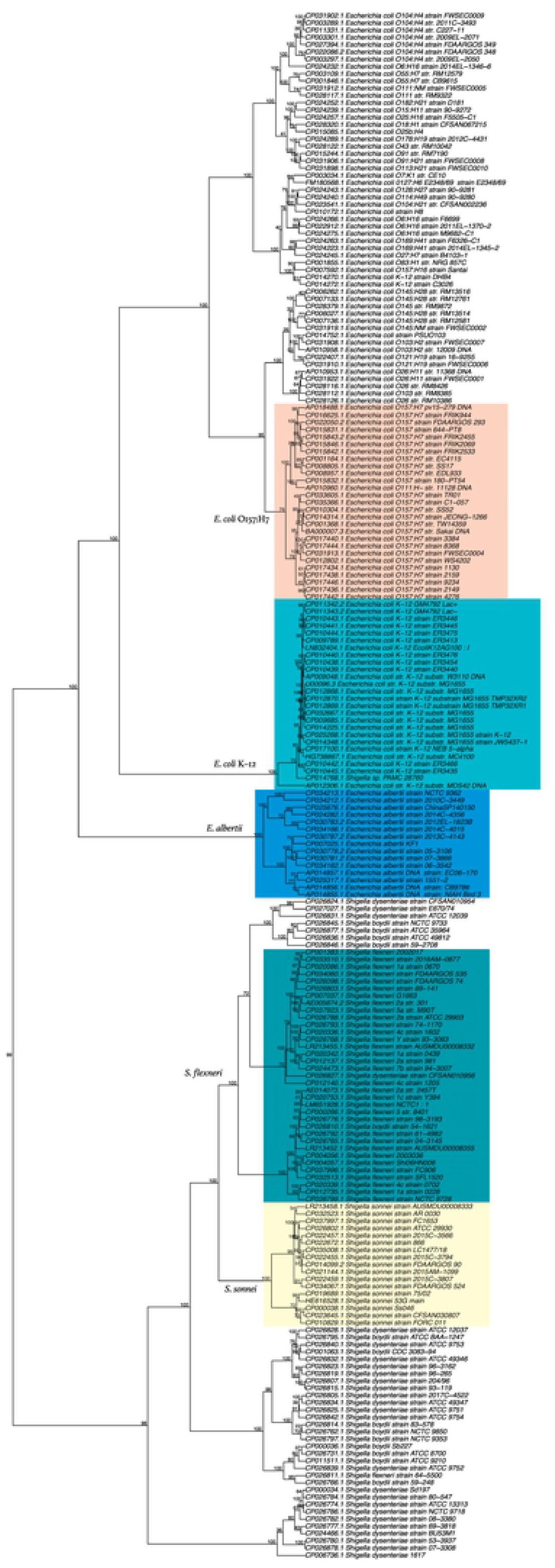
Phylogenetic trees of *E. albertii, Shigella*, and *E. coli* with serotype information based on the pentanucleotide frequencies. The numbers at the nodes indicate the percentage occurrences among 1,000 bootstrap values. Separated groups of species and serotypes are highlighted.

Sims and Kim (2011) reported the whole-genome phylogeny of the *E. coli/Shigella* group based on FFPs [19]. The phylogenetic tree was constructed using all possible features of 24-nucleotide segments, constituting approximately 8.4 million sequences. The compositions of the features were refined to core features of ca 0.56 million with low frequency and low variability. Supplementary Figure 3 suggests that 512 features of pentanucleotides are adequate to construct a phylogenetic tree for *E. coli/Shigella*.

This approach can be applied to any species without defining species-specific gene sets. As shown in Supplementary Figs 3-6, *Campylobacter, Klebsiella*, and *Neisseria* species were clearly distinguished as phylogenetic lineages. In the phylogenetic tree of *Listeria*, serotypes 1/2a, 1/2b, or 1/2c, and 4 were distinguished from each other. The construction of phylogenetic trees using genome sequences takes approximately 10 min. Furthermore, rather than determining the whole genome sequence, this approach only requires clusters of short (100 bp) sequences determined through pyrosequencing.

We provide an application for phylogenetic analysis of species based on the pentanucleotide frequencies as described above. The application Phy5 is written in R and is freely available at github.com/YoshioNakano2021/phy5. The online version of Phy5 can be accessed at https://phy5.shinyapps.io/Phy5R/. FASTA files containing genome nucleotide sequences (one file for one strain) are uploaded onto the site, and Phy5 draws the resultant phylogenetic tree and tables of pentanucleotide frequencies in proportion using aggregated data.

One of us (Y.N.) attempted to create interspecies hybrids among *Ipomoea* species, including some morning glories and used a phylogenetic tree for choice combinations of species. Nucleotide sequences of 30 chloroplast genomes from 24 *Ipomoea* species were downloaded from the GenBank sequence database, and their phylogenetic tree was constructed by the Phy5 system described above (Fig. 5). The chloroplast DNAs were 162–165 kb long, and it took ten seconds to calculate pentanucleotide frequencies and construct the phylogenetic tree. The resultant phylogenetic tree correlated with the trees previously reported, based on a comparison of their specific genes or morphological characteristics [28–30]. In addition, textitI. purpurea, *I. nil, I. tricolor*, and *I. hederacea* are all morning glories, but crossings among *I. purpurea, I. nil*, and *I. hederacea* rarely succeed. Incidentally, *I. quamoclit* and *I. hederifolia* are cypress vines and *I. batatas* is a sweet potato; they cannot successfully be crossed with morning glories.

**Fig 5.**
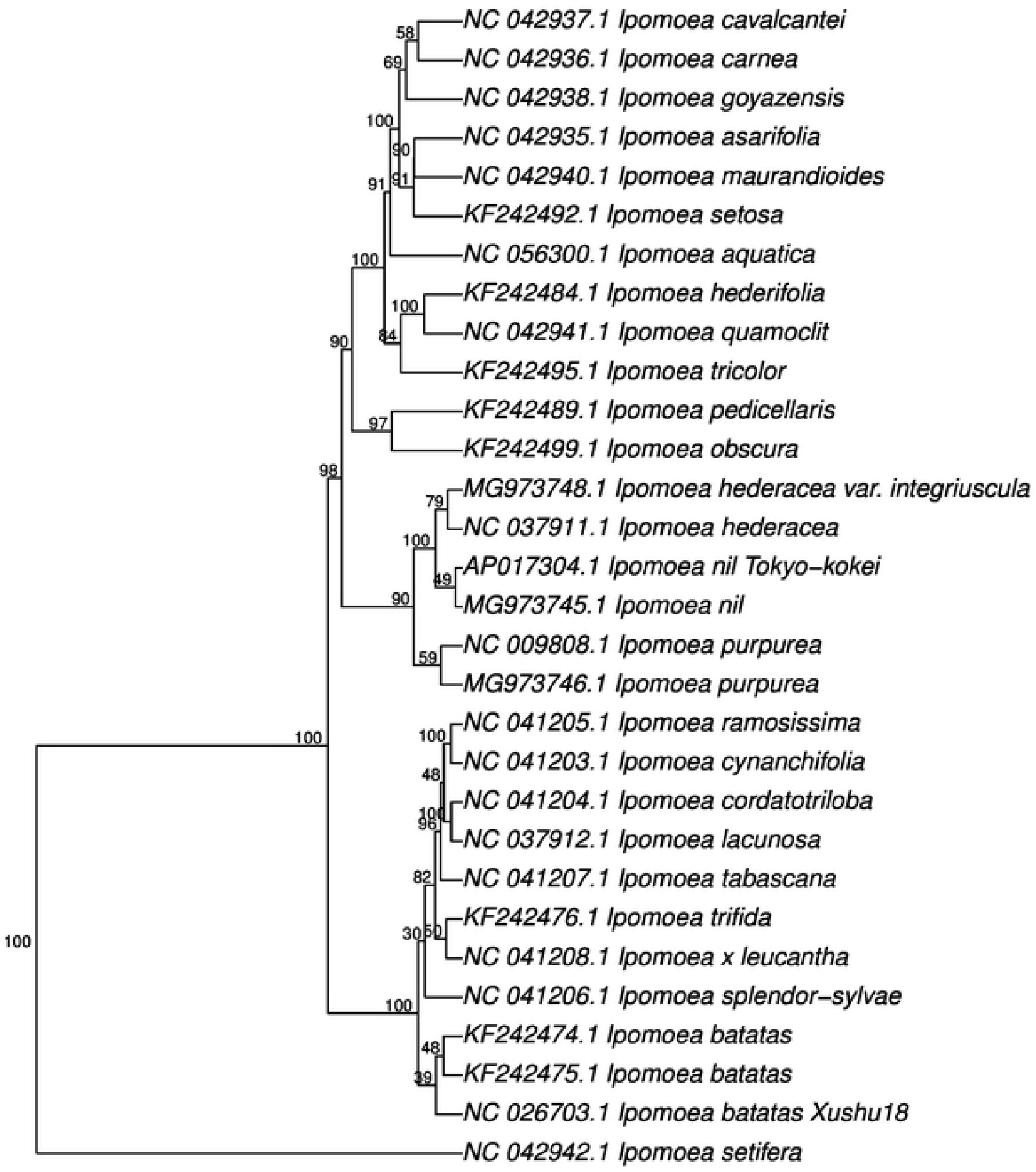
Phylogenetic trees of *Ipomoea* species based on their chloroplast genomes. The numbers at the nodes indicate the percentage occurrences among 1,000 bootstrap values. Separated groups of species and serotypes are highlighted.

As shown above, only closely related species are suitable for this phylogenetic analysis, which is based on pentanucleotide frequencies. Figure 6 shows a phylogenetic tree of various distantly related species, including thermophilic archaea and bacteria. *Pyrococcus* and a *Thermococcus* strain form a clade with the *Streptococcus* species, and these thermophilic archaea are distinct from *Pyrobaculum* and other *Thermococcus* species. The latter thermophilic archaea and thermophilic bacteria, *Thermotoga*, fall within the same clade. Long nucleotide frequencies are suitable for such distant related species, 18 or 24 nucleotides, such as reported by Sims, et al. (2009). [18, 19]. The combination 2^24^ is tedious to calculate, thus a limited sequence set must be selected to increase specificity for the target species.

**Fig 6.**
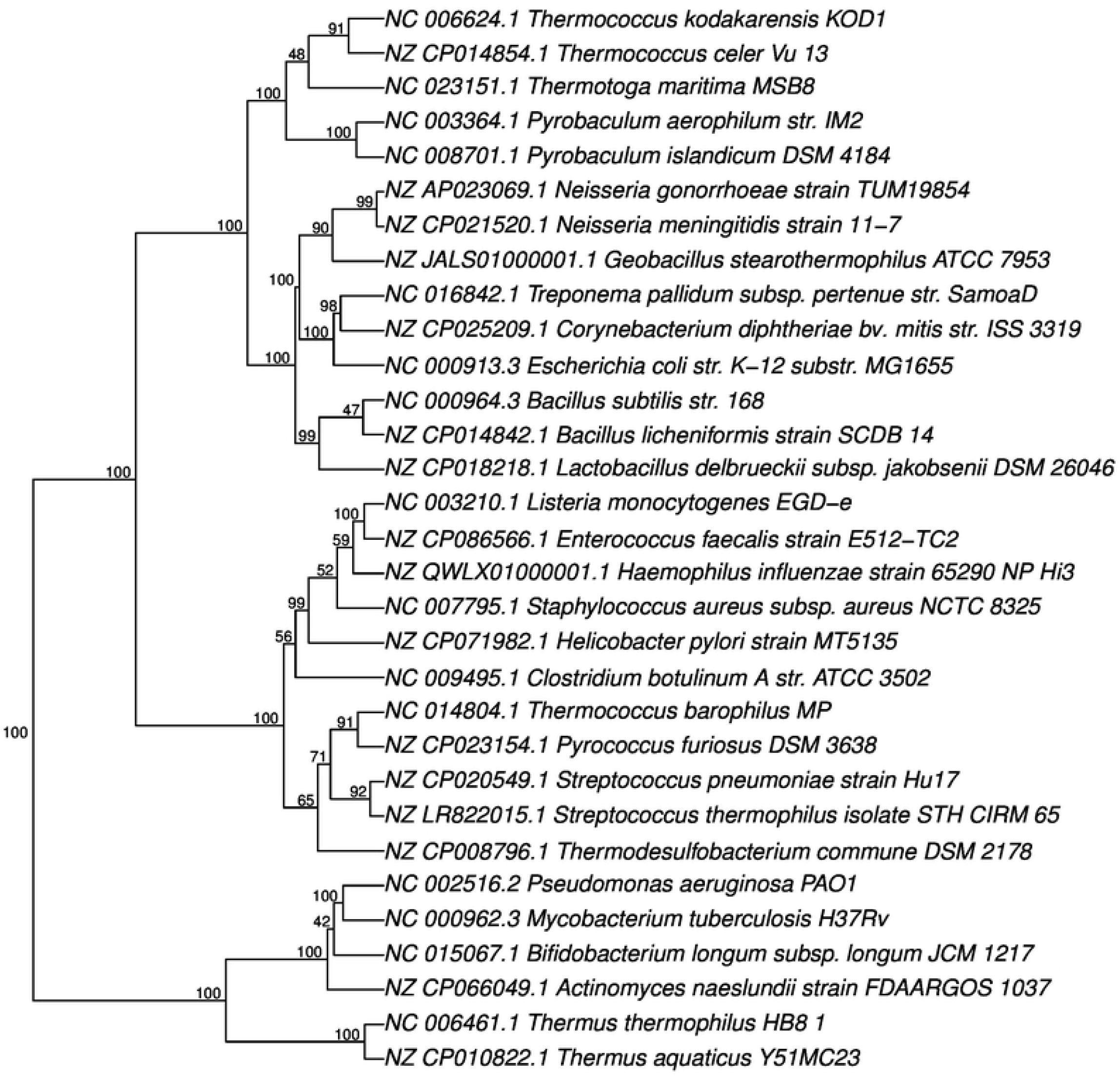
Phylogenetic trees of various distantly related species including archaea and bacteria. The numbers at the nodes indicate the percentage occurrences among 1,000 bootstrap values. Separated groups of species and serotypes are highlighted.

SVM classified 856 strains of *E. coli* and 91 strains of *Shigella*, except one *Shigella* strain with 99.9% accuracy (Table 1).

**Table 1.** Classification of *E. coli* and *Shigella* by SVM.

| Classification | *E. coli* | *Shigella* |
| --- | --- | --- |
| *E. coli* | 856 | 1 |
| *Shigella* | 0 | 91 |

This study shows that phylogenetic analysis based on pentanucleotide profiles offers an alternative for microbial phylogenetic analyses. Future studies are required to identify K-mer sequence combinations that are more specific to bacterial species and develop more efficient calculation methods for microbial phylogenetic analysis.

## Supporting information

**S1 Fig. Phylogenetic trees based on (A) tri-, (B) tetra-, (C) penta-, and (D)hexa-nucleotide frequencies (Supplementary Data 2-4) were constructed**.

The trees were constructed using the Euclidean distance and Ward’s algorithm. The numbers at the nodes indicate the percentage occurrences among 1,000 bootstrap values.

**S2 Fig. Phylogenetic trees for *E. coli/Shigella* based on the pentanucleotide frequencies**. The trees were constructed using the Euclidean distance and Ward’s algorithm. The numbers at the nodes indicate the percentage occurrences among 1,000 bootstrap values.

**S3 Fig. Phylogenetic trees of *Campylobacter* species based on the pentanucleotide frequencies**. The trees were constructed using the Euclidean distance and Ward’s algorithm. The numbers at the nodes indicate the percentage occurrences among 1,000 bootstrap values.

**S4 Fig. Phylogenetic trees of *Klebsiella* species based on the pentanucleotide frequencies**. The trees were constructed using the Euclidean distance and Ward’s algorithm. The numbers at the nodes indicate the percentage occurrences among 1,000 bootstrap values.

**S5 Fig. Phylogenetic trees of *Neisseria* species based on the pentanucleotide frequencies**. The trees were constructed using the Euclidean distance and Ward’s algorithm. The numbers at the nodes indicate the percentage occurrences among 1,000 bootstrap values.

**S6 Fig. Phylogenetic trees of *Escherichia albertii*, including *E. coli* and *Shigella* species based on the pentanucleotide frequencies**. The trees were constructed using the Euclidean distance and Ward’s algorithm. The numbers at the nodes indicate the percentage occurrences among 1,000 bootstrap values.

**S1 File. Supplementary Data 1**. CSV file containing the degenerated pentanucleotide frequencies in the DNA sequences of *Yersinia, E. coli, Shigella, Campylobacter, Klebsiella, Listeria, Neisseria*, and *E. albertii*.

## Acknowledgments

We would like to thank Editage (www.editage.com) for English language editing.

